# Host genetics determines the fate of the *Lactobacillaceae* population in the small intestine

**DOI:** 10.64898/2026.09.25.754556

**Authors:** Helen Beilinson Lietuvninkas, Michael Coyne, Kiera Bass, Madeline Maslyar, Emily Cullum, Jose Villalobos, Alexander Chervonsky, Tatyana Golovkina

## Abstract

Whereas innate pattern recognition receptors bind evolutionary conserved moieties shared by diverse groups of microbes, control over closely related microorganisms is likely based on different principles and can be sensitive to genetic variation of the host. While the influence of host genetics on intestinal microbiome composition has been growing in appreciation, understanding of polymorphic mechanisms that regulate individual bacterial lineages requires complex approaches. Using gnotobiotic mouse monocolonized with *Ligilactobacillus murinus* (ASF361), we found that C57BL/6J (B6) mice harbored significantly more intestinal ASF361 (or other *Lactobacillaceae* genera) than did BALB/cJ (BALB/c) mice. The difference in ASF361 abundance was independent of adaptive immunity and bacteriotoxic factors. In B6 mice, ASF361 occupied ileal crypts, showed greater expression of adhesion-related molecules, and enriched expression of carbohydrate-metabolism pathways, whereas in BALB/c mice, ASF361 remained largely luminal and enriched expressed genes associated with starvation and stress. These findings highlight the importance of host genetic variation in shaping microbial colonization and suggest that such variation should be considered when evaluating probiotic strains and microbiota transplantation.

## Introduction

The intestinal microbiome contributes to host nutrient metabolism, epithelial homeostasis, immune development, colonization resistance, and susceptibility to disease^1^. Some commensals are used as probiotics to improve intestinal health^2^ and some are transplanted to build up colonization resistance to antibiotic-resistant pathogens^3–6^. These applications need to consider the impact of the host’s genetics on the microbes. Microbial community-level changes emerge from both the host’s effects on specific organisms and from interactions among the resident microbes. Thus, defining how the host regulates individual bacterial populations is particularly important.

Studies show that genetically identical individuals (identical twins) harbor more similar microbial communities compared to non-identical twins and that the abundance of some bacterial taxa is heritable^7,8^. Subsequent studies in humans and nonhuman primates have further associated host genetic variation with microbiome composition^7–11^. It was also found that hosts’ genetic polymorphisms were associated with the abundance of some bacterial taxa in mice^12–14^. However, diet, housing, maternal transmission, microbial exposure, and other environmental factors exert a strong influence, complicating efforts to establish casual relationships between host genotype and specific microorganisms^15,16^. Competition for nutrients and spatial niches, metabolic cooperation, and antagonistic interactions may further amplify or obscure a direct host effect^17,18^. Understanding and distinguishing the disparate impact of these possibilities requires experimental systems that can separate host control of an individual bacterium from the influence of the surrounding microbial community.

Gnotobiotic models provide a controlled experimental setting for dissecting these interactions. Reciprocal microbiota transplantation experiments using zebrafish and mouse donors have shown that germfree (GF) recipients impose host-specific selective pressures that reshape introduced microbial communities^19^. In an ex-germfree (exGF) model, the same donor microbiota from specific pathogen-free (SPF) cecal content is introduced into genetically distinct GF recipients, reducing variation arising from maternal transmission, cage effects, and colonization history. Using this approach, we previously showed that host genetics shapes the overall intestinal microbial community^20^ and that some genotype-dependent differences persist despite cross-fostering and cohousing, commonly used to attempt to normalize the microbiome composition between mice^21^. However, the complexity of SPF donor-derived communities makes it difficult to identify the bacteria directly affected by host genotype and the mechanisms responsible.

Defined microbial communities reduce this complexity while preserving biologically relevant host-microbe and interspecies interactions. Here, we used Altered Schaedler’s Flora (ASF), a tractable community of seven bacterial species^22,23^, to identify *Ligilactobacillus murinus* (ASF361) as a commensal whose abundance is directly regulated by host genetics. Furthermore, BALB/cJ (BALB/c) mice monocolonized with ASF361 harbored markedly fewer bacteria than did C57BL/6J (B6) mice and this difference persisted across microbial inputs, vertical transmission, and cohousing. In BALB/c mice, ASF361 was largely excluded from small intestinal crypts and exhibited reduced expression of carbohydrate-metabolism and adhesion pathways. These findings demonstrate that host genetic effects can persist amid other major influences on microbiome composition and must be considered when evaluating bacterial colonization and function.

## Results

### Host genetic variation shapes intestinal *Lactobacillaceae* abundance

We started with a model in which B6 mice harbored similar or greater abundances of cecal *Lactobacillaceae* than BALB/c mice^20^. We repeated experiments in which GF B6 and BALB/c adult mice were gavaged with cecal content from three independent specific pathogen free (SPF) C57BL/6NTac donors to produce exGF mice and analyzed abundance of *Lactobacillaceae* using 16S rRNA sequencing (16S rRNA-seq) (**Figure 1A**). Although the magnitude and statistical significance of the difference varied across donor input communities, *Lactobacillaceae* were generally found to be more abundant in B6 mice.

**Figure 1.**
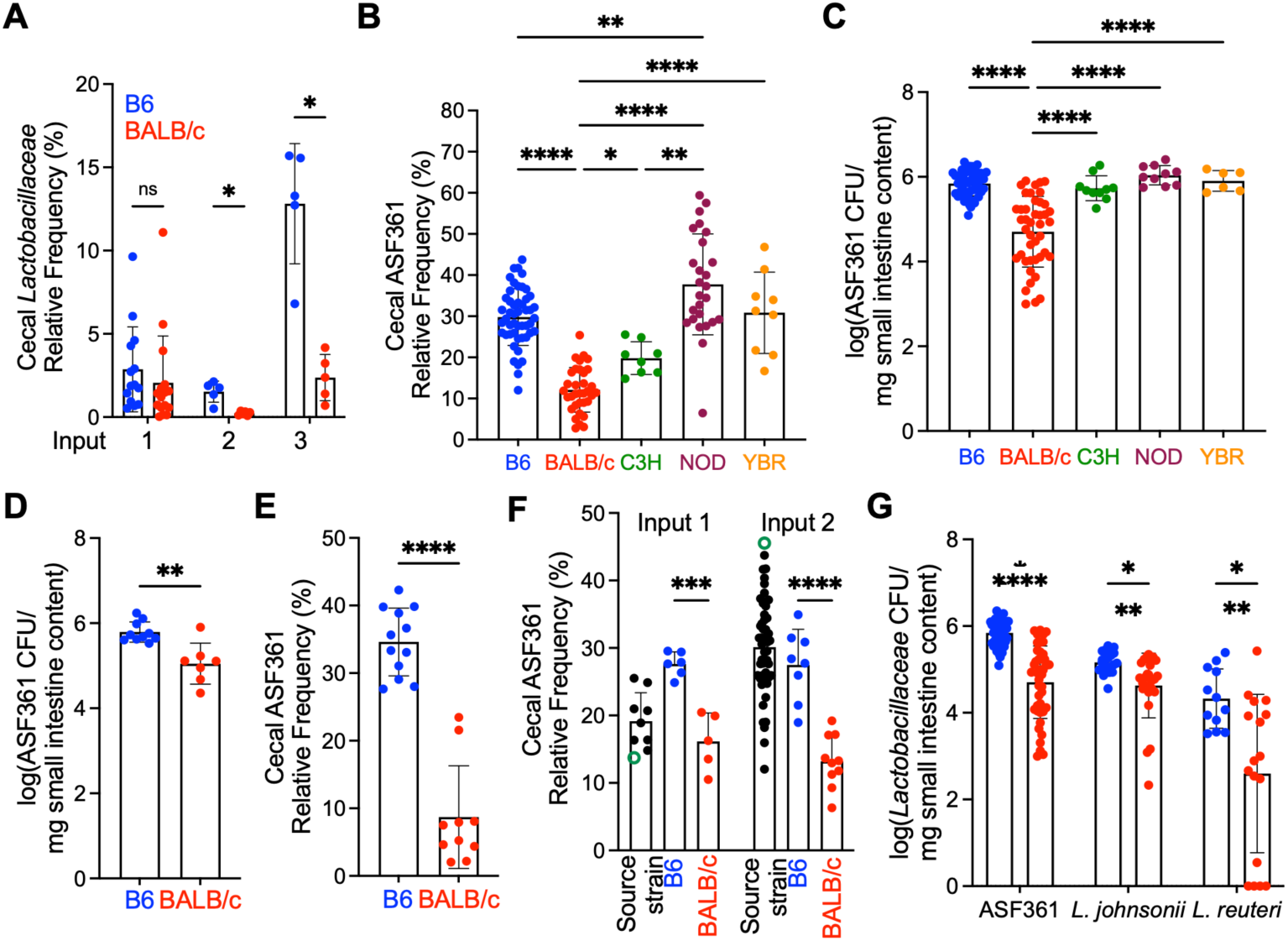
Host genetics shapes *Lactobacillaceae* abundance. (A) Relative frequency of *Lactobacillaceae* taxa in three independent experiments in which GF mice received total cecal contents from C57BL/6NTac mice. (B) Relative frequency of cecal ASF361 in ASF-colonized mice. (C) ASF361 colony-forming units (CFU) in the small intestine of monocolonized mice. (D) ASF361 colony-forming units (CFU) in the small intestine of ASF-colonized mice. (E) Relative frequency of cecal ASF361 in 8-week-old adult progeny of ASF-colonized mice. (F) Relative frequency of ASF361 in the two ASF inocula (green) and in the ceca of their source strains and ASF-colonized recipient mice. Source strain for input 1: C3H; source strain for input 2: B6. (G) *Lactobacillaceae* CFU in the small intestine of monocolonized mice. Unless otherwise indicated, mice were colonized with ASF or monocolonized with *Lactobacillaceae* at 8 weeks of age and analyzed 2 weeks later. BALB/c, BALB/cJ; C3H, C3H/HeN; B6, C57BL/6J; NOD, NOD/ShiLtJ; YBR, YBR/Ei. Relative frequency determined by 16S rRNA-seq. Each dot represents an individual mouse. Data were pooled from 2-9 independent experiments. Statistical significance was assessed using two-tailed unpaired Student’s *t*-test (A, D-G) or two-way ANOVA (B, C). Data are presented as mean ± SD. \**P* < 0.05, \*\**P* < 0.01, \*\*\**P* < 0.001, \*\*\*\**P* < 0.0001; ns, not significant.

To reduce community complexity, we turned to Altered Schaedler’s Flora (ASF), a defined murine community that includes *Ligilactobacillus murinus*, ASF361^22,23^. Adult GF mice from several inbred strains were colonized with ASF, and their cecal microbiomes were analyzed by 16S rRNA-seq two weeks later (**Figures 1B, S1A-B**). ASF361 abundance was statistically higher in B6, NOD/ShiLtJ (NOD), and YBR/Ei (YBR) mice than in BALB/c, with a smaller difference observed in C3H/HeN (C3H) mice.

It was possible that the lower abundance of ASF361 in BALB/c mice was affected by interactions with other ASF members. Thus, we monocolonized GF mice from various strains with ASF361 and quantified colony-forming units (CFU) in the small intestine (**Figure 1C**). Although *Lactobacillaceae* colonize the entire intestinal tract, their tolerance of acidic, bile-rich, and relatively oxygenated conditions makes them particularly well adapted to the small intestine^24^. Gnotobiotic BALB/c mice harbored fewer ASF361 CFU than every other mouse strain tested, including B6 mice (**Figure 1C**) consistent with 16S rRNA-seq data obtained with ASF-colonized mice (**Figure 1B**) and with ASF361 CFU in the small intestine of ASF-colonized mice (**Figure 1D**). We focused subsequent analyses on B6 and BALB/c mice because genetically modified strains of these backgrounds were more readily available in GF settings.

To test whether natural (during birth and early feeding) colonization would produce similar results, we bred ASF-colonized mice and analyzed the cecal microbiomes of their adult offspring (**Figures 1E, S1C**). Mice colonized from birth retained the strain-specific difference, with ASF361 remaining more abundant in B6 than in BALB/c mice.

ASF is stably maintained and propagated in gnotobiotic mice. Since input microbiota composition influences the composition of the resulting output community^20^, we colonized GF B6 and BALB/c mice with ASF communities from two different source mouse strains that differed in ASF361 abundance, B6 and C3H mice (**Figures 1F, S1D**). ASF361 was once again found to be more abundant in B6 than in BALB/c recipients, regardless of its relative abundance in the input community.

We then tested whether the factor(s) controlling ASF361 abundance could be horizontally transferred by cohousing monocolonized B6 and BALB/c mice either with mice of the same strain or with mice of the other strain (**Figure S1E**). Regardless of housing conditions, B6 mice continued to harbor more ASF361 in the small intestine than did BALB/c mice. Together, these data indicate that a host-intrinsic factor limits ASF361 abundance in BALB/c mice regardless of the age at colonization and that cohousing does not overcome this phenotype. This factor also acts rapidly, as BALB/c mice colonized with ASF for just one week harbored fewer ASF361 than B6 mice (**Figure S1F**).

Finally, we asked whether this phenotype was specific to ASF361 or extended to other members of the *Lactobacillaceae* family by colonizing GF B6 and BALB/c mice with *L. johnsonii* or *L. reuteri*, two additional murine commensals (**Figure 1G**). B6 mice harbored more of both species than BALB/c mice, indicating that this strain-dependent difference extends across multiple *Lactobacillaceae* genera.

### Suppression of ASF361 in BALB/c mice is independent of bacteriotoxic factors and adaptive immunity

The intestinal mucosa acts as a nutrient source for *Lactobacillaceae* but also contains numerous innate antimicrobial molecules and peptides^25^. To determine whether BALB/c mucus lacks a nutrient source or contains a strain-specific bacteriotoxic factor suppressing ASF361, we cultured in small intestine mucus from GF B6 and BALB/c mice and measured growth over 24 hours (**Figure 2A**). Growth did not differ, providing no evidence that BALB/c mucus is intrinsically toxic to ASF361 nor does it lack nutritional capacity *in vitro*. α-Defensins (Defa) produced by Paneth cells in the crypts of Lieberkühn^26^ are major polymorphic antimicrobial peptides between B6 and BALB/c mice^20^, but have a very short half-life and may not be captured in our *ex vivo* culture. To test if a BALB/c-specific Defa affects ASF361, we monocolonized BALB/c mice deficient in matrix metalloproteinase-7 (MMP7), which converts inactive pro-defensins into their active forms^27^. Small intestine bacterial burdens did not differ between WT and MMP7-deficient mice, indicating that Defa do not account for the strain-dependent phenotype (**Figure 2B**).

**Figure 2.**
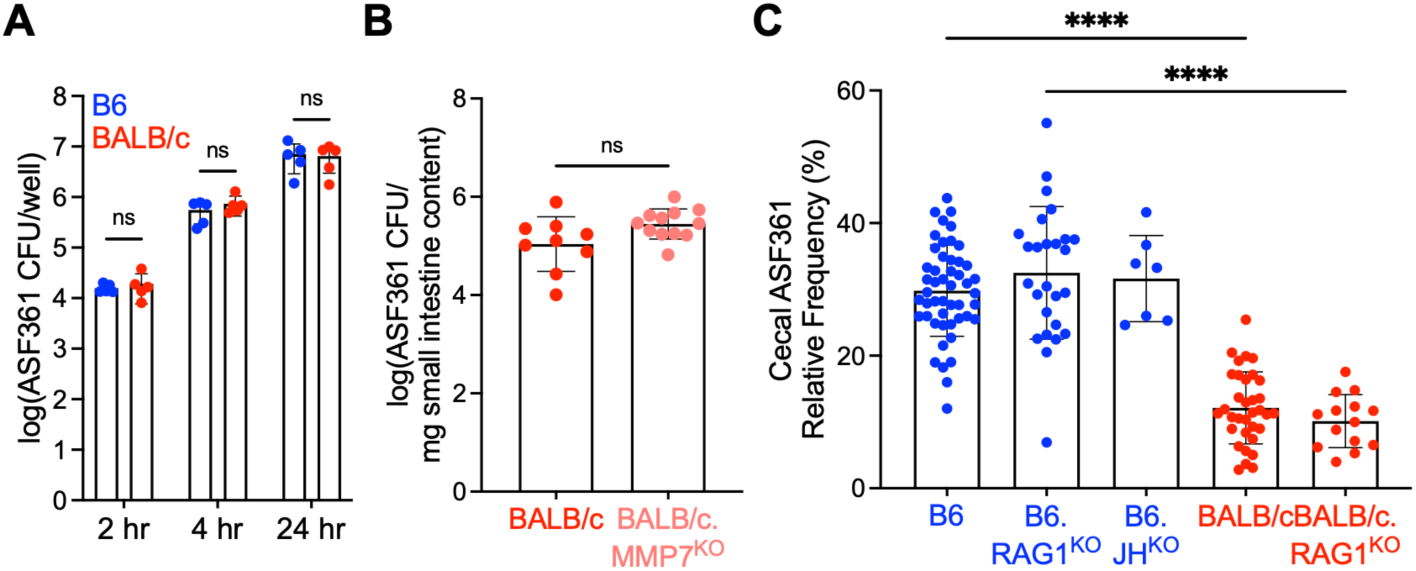
Suppression of ASF361 in BALB/c mice is independent of bacteriotoxic factors and adaptive immunity. (A) ASF361 colony-forming units (CFU) after growth on small intestine mucus from germ-free mice. Each dot represents the mean bacterial growth across three replicate wells containing mucus from an individual mouse. (B) ASF361 colony-forming units (CFU) in the small intestine of monocolonized WT and MMP7-deficient mice. (C) Relative frequency of cecal ASF361 in ASF-colonized RAG1-and JH-deficient mice. Relative frequency determined by 16S rRNA-seq. Mice were colonized with ASF or monocolonized with ASF361 at 8 weeks of age and analyzed 2 weeks later. Each dot represents an individual mouse. Data were pooled from 1-9 independent experiments. Statistical significance was assessed using a two-tailed unpaired Student’s *t*-test (A-C). Data are presented as mean ± SD. \*\*\*\**P* < 0.0001; ns, not significant.

We then tested whether adaptive immunity contributes to ASF361 suppression in BALB/c mice. RAG1-deficient mice, which lack T and B cells, harbor microbial communities distinct from those of immunocompetent mice, demonstrating that adaptive immunity can contribute to shaping of the intestinal microbiome ^20,28,29^. We therefore colonized GF RAG1-deficient B6 and BALB/c mice, as well as GF JH-deficient B6 mice that lack B cells, with ASF (**Figures 2C, S2**). ASF361 abundance in RAG1-, JH-deficient and wild-type (WT) mice were similar. Thus, adaptive immunity is not required for BALB/c-associated control of ASF361.

### ASF361 is excluded from BALB/c ileal crypts

We next examined the distribution of ASF361 in the small intestine by Gram-staining ileal sections from B6 and BALB/c mice after two weeks of monocolonization of GF RAG1-deficient mice (**Figure 3A, S3A**). RAG1-deficient mice were used as they phenocopy WT counterparts. In B6 mice, ASF361 penetrated the intervillous spaces and occupied the crypts, whereas in BALB/c mice, it remained largely restricted to the intestinal lumen. Consequently, nearly all ileal crypts in B6 mice contained ASF361, compared to only one-quarter of those in BALB/c mice (**Figure 3B, S3B**). The extensive penetration of ASF361 into B6 intervillous spaces suggests that B6 mice possess a factor that promotes colonization or that BALB/c mice possess a factor that suppresses it. The suppressive effect of such a factor may depend on the intestinal environment and therefore was not apparent in our *in vitro* experiments.

**Figure 3.**
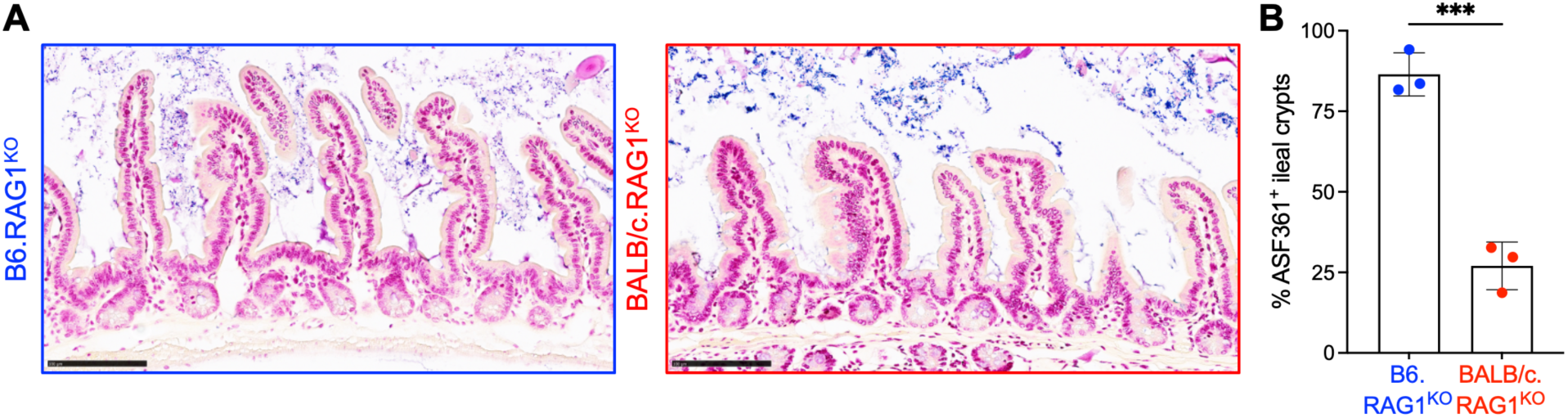
ASF361 is excluded from BALB/c ileal crypts. (A) Gram staining of Carnoy’s-fixed ileal sections from mice monocolonized with ASF361. Each bar is 100 μm. (B) Percentage of ileal crypts containing ASF361. Each dot represents an individual mouse. Locations of analyzed crypts are shown in Figure S3B. Mice were colonized with ASF361 at 8 weeks of age and analyzed 2 weeks later. Statistical significance was assessed using a two-tailed unpaired Student’s *t*-test. Data are presented as mean ± SD. \*\*\**P* < 0.001.

### Carbohydrate metabolism is reduced in ASF361 isolated from BALB/c mice

We reasoned that a (B6 x BALB/c)F1 cross should help us to distinguish between these two possibilities. Thus, we crossed GF WT or GF RAG1-deficient B6 and BALB/c mice and colonized their F1 offspring with either ASF361 alone or with the entire ASF consortium (**Figures 4A, S4A-B**). Monocolonized F1 mice exhibited an intermediate bacterial burden relative to the two parental strains (**Figure 4A**) indicating that neither parental phenotype is fully dominant or recessive. However, in ASF-colonized F1 mice, the relative abundance of cecal ASF361 resembled that observed in BALB/c mice (**Figures S4A-B**), indicating that other bacteria in this community may contribute to the overall phenotype dictated by the host genetic factors.

**Figure 4.**
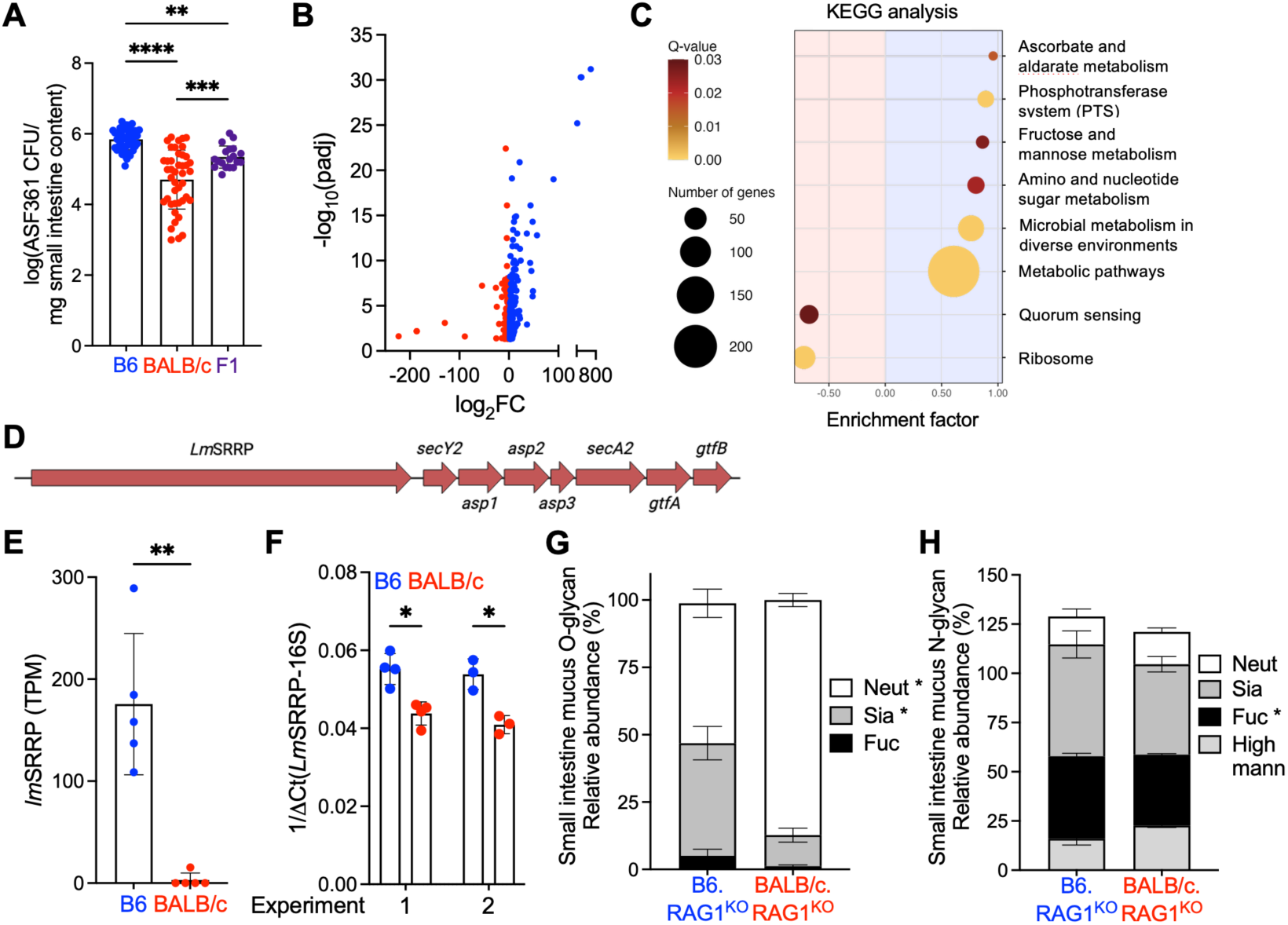
Carbohydrate metabolism is reduced in ASF361 isolated from BALB/c mice. (A) Small intestine ASF361 CFU in monocolonized mice. (B) Volcano plot of differentially expressed genes (DEGs) in ASF361 isolated from monocolonized B6 or BALB/c mice. Genes enriched in bacteria from B6 and BALB/c mice are shown in blue and red, respectively; each dot represents one gene. (C) KEGG pathway enrichment analysis of the DEGs identified in (B). (D) Schematic of SecA2/Y2 locus in ASF361. (E) Transcripts per million (TPM) for *Lm*SRRP from ASF361 isolated from monocolonized B6 or BALB/c mice, analyzed by RNA-seq. (F) qPCR analysis of *Lm*SRRP expression from ASF361 from two independent monocolonization experiments. Relative abundances of (G) O-and (H) N-glycans in small intestine mucus from GF mice. Fuc, fucosylated; Sia, sialylated; Neut, neutral complex; High mann, high mannose. Each dot represents an individual mouse. Mice were colonized with ASF361 at 8 weeks of age and analyzed 2 weeks later. Statistical significance was assessed using two-tailed unpaired Student’s *t*-test (A, F-H), edgeR (B, E), and the Benjamini–Hochberg method (C). Data are presented as mean ± SD. \**P* < 0.05, \*\**P* < 0.01, \*\*\**P* < 0.001, \*\*\*\**P* < 0.0001; ns, not significant.

To get mechanistic insight into our observations, we examined the gene expression by ASF361 in the distinct small intestinal environments. RNA sequencing (RNA-seq) of bacteria isolated from monocolonized mice identified 344 differentially expressed genes (DEG) between the two groups (**Figure 4B**). Genes expressed at higher levels in ASF361 from B6 mice were enriched in metabolic pathways including carbohydrate-uptake, whereas those expressed at higher levels in bacteria from BALB/c mice were enriched in quorum-sensing and ribosomal pathways (**Figure 4C**).

Because ASF361 associates closely with the gastrointestinal epithelium and forms biofilm-like structures, mucosal attachment contributes to its persistence^30^. Limited access to adhesion sites or lack of adhesin expression could reduce bacterial retention and restrict access to locally available nutrients, diminishing metabolism pathways in the bacteria. Given the established role of mucosal attachment in the colonization and persistence of *Lactobacillaceae* genera^31–33^, we hypothesized that ASF361 might express adhesion-related genes less efficiently in the BALB/c small intestine. In *L. reuteri*, a serine-rich repeat protein (SRRP) adhesin with a long signal sequence containing a KxYKxGKxW motif is encoded by the SecA2/Y2 accessory secretion locus and contributes to epithelial adhesion^34–36^. Sequencing the ASF361 genome revealed an SRRP-encoding homolog within the SecA2/Y2 locus, which we designated *Lm*SRRP (**Figure 4D**). *Lm*SRRP expression was higher in bacteria recovered from B6 mice than in those recovered from BALB/c mice, as shown by RNA-seq (**Figure 4E**) and confirmed by qPCR in two additional independent monocolonization experiments (**Figure 4F**). Importantly, the difference was not reproducible after *ex vivo* culturing in mucus isolated from GF B6 and BALB/c mice (**Figure S4C**).

These findings support a model in which genetically determined features influence ASF361 adhesin expression and access to crypt-associated adhesion sites. BALB/c mice likely encode an allele that fails to stimulate or actively suppresses expression of the host substrate required for bacterial adhesion. The expression of several SecA2/Y2-dependent substrates responds to metabolic or environmental conditions^37,38^, with host glycans being plausible mediators of the strain-dependent phenotype. In addition to serving as metabolic substrates and environmental signals, host glycans can also function as adhesin receptors^36,39^. Thus, the reduced *Lm*SRRP expression and crypt occupancy by ASF361 in BALB/c mice could reflect limited availability of a cognate receptor or the altered presentation of a glycan that regulates bacterial gene expression. O-and N-linked glycan analysis of GF mucus revealed extensive differences between the small intestinal glycan profiles of B6 and BALB/c mice (**Figure 4G-H**). Potential candidate glycans driving ASF361 suppression in BALB/c mice are likely present within this pool.

## Discussion

Although host genetics is widely recognized as a determinant of microbiome composition^7–9,12–14,20^, the mechanisms of host polymorphic genes regulation of individual bacterial lineages remain poorly understood. Here, we used a gnotobiotic approach to show how host genetics influences the abundance, localization, and transcriptional state of ASF361. Introducing identical communities or single bacterial species into genetically distinct GF hosts allowed us to separate direct host effects from those that emerged in the presence of other community members. This framework can be extended to identify polymorphic host factors that regulate specific commensal populations.

The effect of host genotype was evident across several experimental conditions. BALB/c mice consistently harbored fewer ASF361 than B6 mice following monocolonization, colonization with ASF, vertical transmission, and cohousing. The phenotype also extended to *L. johnsonii* and *L. reuteri*, suggesting that the responsible host feature affects multiple *Lactobacillaceae* genera. In addition, mucus isolated from the BALB/c small intestine did not inhibit ASF361 growth relative to growth on B6 mucus. Thus, the phenotype was not explained by inherent bacteriotoxic or nutritional properties of the mucus. Instead, our results support a model in which the BALB/c intestinal environment does not allow ASF361 access to the crypt niche and is associated with reduced expression of an adhesin. Adhesion may allow ASF361 to persist near the epithelium and access spatially restricted nutrients or host signals^30,34,36^.

Our findings also distinguish bacterial presence from bacterial physiological state. Although the same strain of ASF361 colonized both B6 and BALB/c mice, bacteria recovered from the two strains occupied different anatomical niches and exhibited markedly different transcriptional profiles. In B6 mice, ASF361 entered ileal crypts and showed greater expression of carbohydrate-metabolism pathways and adhesin expression. In BALB/c mice, the bacteria remained predominantly luminal and expressed genes associated with starvation or stress-related responses. Because ASF361 grew comparably *ex vivo* in mucus isolated from either GF B6 or from BALB/c mice, the strain-dependent differences in ASF361 metabolism and adhesin expression likely reflect colonization-induced changes in host mucosal glycosylation^39^ or other epithelium-associated cues. Host-derived glycans can induce adhesive phenotypes in several Gram-positive bacteria, including *Lactobacillaceae*. For example, expression of MbpA, a MUC5B-binding adhesin in *Streptococcus gordonii*, depends on MUC5B^40^, exposure to porcine mucin induces the surface localization of several moonlighting adhesins, including EF-Tu in *L. delbrueckii* and *L. plantarum*^41^ and pyruvate kinase and fructose-bisphosphate aldolase in *L. acidophilus*^42^. Synthetic polymers bearing β-galactose-or β-lactose increase the adhesion of *L. fermentum* to a mucin-coated surface^43^. Moreover, it has been shown that Gram-positive adhesins recognize host sialylated glycans^44,45^. Together, these observations suggest that polymorphic host factors could restrict ASF361 colonization in BALB/c mice at two levels: by failing to induce or actively inhibiting *LmSRRP* expression and by providing fewer sialylated adhesion sites compared to the number of these sites available in B6 mice.

Interestingly, the F1 experiments revealed that both host and microbial factors can produce amplified signals for colonization with a particular bacterial lineage. Monocolonized F1 mice had an intermediate ASF361 burden, whereas ASF-colonized F1 mice exhibited a stronger BALB/c-like phenotype. Whereas in the former case, the result can be interpreted as a dilution of the positive cue (from B6 parent), in the latter case, the BALB/c influence appears dominant. This dominance, however, more likely results from suppressive influence of other ASF members that are not present during monocolonization. These results are consistent with two previous findings. First, comparisons of B6 and BALB/c mice with their reciprocal F1 crosses (B6 x BALB/c and BALB/c x B6) showed that host genetic background influenced microbiome composition more strongly than maternal microbiome composition or cohousing with another strain^46^. Second, SPF F1 mice from both reciprocal F1 crosses had *L. johnsonii* abundances intermediate between those of the parental strains^47^. Competition for nutrients or spatial niches could make an intermediate host environment less permissive in the presence of other bacteria. Specific interspecies interactions, including the reported *in vitro* parasitic relationship between ASF361 and ASF519^48^, could also contribute. These findings indicate that other bacteria can exacerbate a colonization disadvantage established by host genetics.

Our findings do not diminish the importance of environmental factors in shaping commensal populations^15,16^; instead, they show that housing, microbial transmission, and interspecies interactions operate within the ecological conditions that are established by host genetics. BALB/c mice supported lower ASF361 burdens and altered the bacterium’s physiological state even during monocolonization in the absence of completion from other bacteria. Host genetics can therefore impose persistent constraints on the abundance, localization, and physiological state of an intestinal commensal. Because dietary interventions, fecal microbiota transplants, and probiotics produce variable responses among individuals^49–51^, and host genotype contributes to this variation^52–54^, host compatibility must be considered when interpreting microbiome variation and predicting the outcomes of microbiome-directed clinical interventions.

## Methods

The studies described here were reviewed and approved by the Animal Care and Use Committee at The University of Chicago accredited by the Association for Assessment and Accreditation of Laboratory Animal Use (AAALAC).

### Mouse lines

#### Germ-free (GF) mice

GF C57BL/6J (WT, JH^KO^, and RAG1^KO^), BALB/cJ (WT, RAG1^KO^, and MMP7^KO^), C3H/HeN, NOD/ShiLtJ non-obese diabetic, and YBR/Ei mice were re-derived at Taconic Biosciences through an embryo transfer procedure. GF (C57BL/6J x BALB/cJ)F1 and (C57BL/6J.RAG1^KO^ x BALB/cJ.RAG1^KO^)F1.RAG1^KO^ animals were generated by crossing GF animals. GF animals were maintained in our gnotobiotic mouse colony at The University of Chicago. Mice were eight weeks old at the start of experiments unless otherwise indicated and an equal mix of males and females were used as no sex-based difference in phenotype was observed (data not shown).

### Monitoring sterility in gnotobiotic isolators

DNA was extracted from freshly frozen fecal pellets using a bead-beating/phenol-chloroform extraction protocol and amplified with a set of primers that hybridize to all bacterial 16S rRNA gene sequences^55^. Weekly tests were conducted using fecal samples from individual cages. Microbiological cultures were set up with GF fecal pellets, positive (SPF) fecal pellets, sham (sterile saline), and negative (sterile culture medium) controls for each sample batch. Samples were inoculated into BHI, Nutrient, and Sabaroud Broth tubes. Each sample was incubated at 37°C and 42°C in aerobic and anaerobic environments. Cultures were followed for five days and monitored daily for evidence of growth.

### Generation of BALB/cJ.MMP7^KO^ mice

For the generation of BALB/cJ.MMP7^−/−^ mice, two guide RNAs were used to target exons 2 (5’-TAGTCTAGTGGACAACCTCA-3’) and 4 (5’-ACTTTGACAAGGATGAGTAC-3’). Founder mice and their offspring were genotyped using primers in intron 1-2 (5’-CCCCAACCTTTCTTGTGCTTG-3’) and intron 4-5 (5’-GGCTAGGTAGGGGCAGCTAA-3’). A line from a single founder with a 940 bp deletion was developed and the founder was crossed to WT BALB/cJ mice for two generations. Heterozygous mice were then intercrossed to produce a homozygous knockout strain used in subsequent studies.

#### Bacteria and consortia

##### Altered Schaedler’s Flora (ASF)

ASF was propagated in gnotobiotic mice in our gnotobiotic colony, as it a nonculturable consortia. All but one species (ASF360, *Lactobacillus acidophilus*) was found to be maintained in our ASF colony. Donors were euthanized and their cecal contents were removed and snap frozen at −80°C.

### Lactobacillaceae strains

ASF361 was previous isolated from ASF by plating suspended fecal matter from ASF colonized mice onto de Man, Rogosa, and Sharpe (MRS) agar^56^, a selective media for *Lactobacillaceae*. *L. johnsonii* were isolated from an SPF WT B6 female mouse from the University of Chicago. *L. reuteri* (ATCC PTA 6475) was a kind gift from Dr. Erika Claud. All *Lactobacillus* strains were grown on MRS agar or in MRS broth at 37°C anaerobically using AnaeroPack jars and GasPaks.

The genome of ASF361 used in these studies was sequenced by hybrid bacterial genome sequencing performed by Plasmidsaurus. Genomic DNA extraction was done using the ZymoBIOMICS DNA Miniprep Kit (Zymo Research, D4300). Long-read sequencing using Oxford Nanopore Technologies chemistry (Oxford Nanopore, R10.4.1) and short-read Illumina paired-end (2×150 bp) sequencing were integrated via the Plasmidsaurus hybrid workflow and polished using Polypolish (v0.6).

#### Colonization of GF mice

All mice, unless noted, were eight-week-old GF miceat the time of colonization with equal numbers of males and females used. Monocolonization: *Lactobacillaceae* strains were introduced to recipient GF mice by oral gavage (200 μL/mouse) of overnight liquid culture grown anaerobically at 37°C from single colonies diluted 1:100 in sterile 1x PBS. ASF: Frozen cecal contents were immediately resuspended in sterile 1x PBS and gavaged (200 μL/mouse) into recipient GF mice. Total cecal: SPF cecal contents used for colonization were obtained from C57BL/6NTac mice purchased from Taconic Farms. Donors of cecal contents were euthanized, and cecal content was removed and snap frozen at −80°C. Contents were prepared the same was as ASF for gavage.

#### DNA extraction and 16S rRNA amplicon sequencing (16S rRNA-seq)

For most experiments, cecal samples were collected into autoclaved Eppendorf tubes and snap frozen at −80°C. DNA extraction from cecal samples was performed using QIAamp PowerFecal Pro DNA Kit (Qiagen, Germantown, Maryland). Barcoded dual-index universal bacterial primers 563F and 926R were used to PCR-amplify the V4-V5 hypervariable region of the 16S rRNA gene. Illumina compatible libraries were generated using the Qiagen QIASeq 1-step amplicon kit and sequencing was performed on the Illumina MiSeq platform using 2×250 paired-end reads.

Regardless of sequencing methodology, raw gene sequence data was demultiplexed and processed through the DADA2 pipeline in R Studio (version 2025.09.2+418)^57^. Amplicon sequence variants (ASVs) were identified, and taxonomy was assigned to the family or genus level using the Silva NR99 database^58^. For exGF mice colonized with total cecal contents, analyses were conducted using the relative abundance of bacterial genera in each sample. For ASF colonized mice, analyses were performed using the relative abundance of bacterial species.

#### Statistical analysis

Statistical analyses of 16S rRNA gene sequencing data were performed in RStudio. Details of the statistical analyses for each experiment, including statistical tests used and the exact value of *n* (where *n* represents the number of animals), are provided in the corresponding figure legends. Randomization and blinding were used for all experiments where possible.

#### Data and code availability

All datasets and sequencing reported in this study will be deposited in public repositories before peer-reviewed publication and are available from the corresponding authors upon reasonable request.

### Enumeration of Lactobacillaceae in gastrointestinal contents

Total small intestine contents, including both luminal contents and mucus, were scraped and collected into sterile tubes, weighed, and resuspended in sterile 1x phosphate-buffered saline (PBS). Samples were homogenized thoroughly and 10-fold serial dilutions were prepared in sterile 1x PBS. Aliquots (triplicates of 5 μL) were plated onto de Man, Rogosa, and Sharpe (MRS) agar plates. Plates were incubated anaerobically at 37°C overnight using an anaerobic GasPak system. Following incubation, bacterial colonies were enumerated and colony-forming units (CFU) were calculated based on dilution factor. Bacterial burden was expressed as CFU per milligram of total small intestine contents (log(CFU/mg)).

#### Gram-staining of ilea of monocolonized mice

Ileal tissues were fixed in Carnoy’s fixative for 1 hr on ice and transferred to 70% ethanol, embedded in paraffin, and sectioned at 5 μm. Tissue sections were deparaffinized in xylene, rehydrated through a graded ethanol series to distilled water, and subjected to Gram staining using a standard Gram staining protocol. Briefly, sections were stained with crystal violet, treated with Gram’s iodine, decolored with acid alcohol, and counterstained with safranin. Stained sections were imaged by brightfield microscopy. Histological sections were imaged using a NanoZoomer S360 whole-slide scanner (Hamamatsu Photonics). To enumerate crypt occupancy, two continuous regions from the ilea and jejunum were randomly selected and crypts were evaluated for presence or absence of bacteria.

#### RNA extraction and sequencing from ASF361 from monocolonized mice

RNA from total small intestine content (lumen and mucus) of C57BL/6J and BALB/cJ monocolonized with ASF361 for two weeks was isolated using the guanidinium thiocyanate/cesium chloride gradient method as described^59^. Oligo-dT based polyA-plus RNA depletion was performed to eliminate host-derived mRNA. rRNA was depleted using a Ribo-Zero rRNA removal kit (Illumina) and RNA sequencing was performed (2×150 bp paired end) using the Illumina NovaSeq 6000 platform.

#### RNA-seq analysis and pathway analysis

After adapter and quality trimming of all reads using utilities included in the BBMap package of bioinformatics tools (v.38.90), the reads were mapped using the Bowtie2 short-read aligner (v.2.4.2)^60^ to the KEGG-annotated *L. murinus* CR147 reference genome (assembly ASM328811v1; BioProject PRJNA411912). Alignment files were converted to sorted and indexed BAM files using SAMtools (v.1.11)^61^ and BEDtools (v.2.30.0)^62^ was used to compare the alignments with a General Feature Format (GFF) file containing protein-coding gene intervals. Differential gene expression was analyzed independently using DESeq2 (v.1.30.0)^63^ and edgeR (v.3.32.1)^64^. Differentially expressed genes (DEGs) were defined as those with fold-change (FC) values of ≥|2| and if the adjusted *P* value (padj for DESeq2 and false discovery rate (FDR) for edgeR) was ≤0.05.

Gene set enrichment analysis was performed using clusterProfiler (v.4.18.4)^65^. Genes were ranked by the DESeq2 Wald statistic and enrichment was evaluated against Kyoto Encyclopedia of Genes and Genomes (KEGG) pathway annotations^66^. *P* values were adjusted for multiple testing using the Benjamini-Hochberg method, and pathways with an adjusted *P* value of ≤0.05 were considered significantly enriched.

The CR147 reference was used because it has organism-specific gene identifies linked directly to KEGG pathway annotations, whereas the Plasmidsaurus hybrid assembly of our experimental strain did not have corresponding KEGG annotations. Although CR147 was not the exact experimental strain, reciprocal BLAST analysis of CR147 and our assembly identified corresponding sequences for most genes, indicating a high degree of similarity between the two genomes (data not shown). We therefore used the CR147-based analysis to evaluate broad pathway-level changes, while recognizing that it may not capture all strain-specific, plasmid-associated, duplicated, or highly divergent genes. The hybrid assembly was used to resolve strain-specific genomic features, including the SRRP putative adhesin in the SecA1/Y2 operon.

#### Reverse transcription and quantitative PCR

Following RNA extraction, contaminating genomic DNA was removed using the DNA-free DNA Removal Kit (Invitrogen) according to the manufacturer’s instructions. cDNA was synthesized from DNase-treated RNA using SuperScript IV Reverse Transcriptase (Invitrogen) and a random hexamers according to the manufacturer’s instructions.

Quantitative PCR (qPCR) was performed using iTaq Universal SYBR Green Supermix (Bio-Rad Laboratories) on an Applied Biosystems QuantStudio 3 Real-Time PCR System. Reactions contained 0.1 ng cDNA and 0.5 μM of each primer in a final volume of 10 μL. Amplification consisted of polymerase activation at 95°C for 30 s, followed by 40 cycles of 95°C for 5 seconds and 60°C for 30 seconds. *LmSRRP* expression was normalized to bacterial 16S rRNA and relative expression was calculated as 1/(Ct(*LmSRRP*)-Ct(16S)). Primer sequences were: *LmSRRP* F (5’-TTGTTCCAGGCCATTACCCC-3’); *LmSRRP* R (5’-TCAAGGCTCAAACACTTCGGA-3’); 341F (5’-CCTACGGGAGGCAGCAG-3’); 519R (5’-GWATTACCGCGGCKGCTG-3’). Original sequences of 16S rRNA primers 341F and 519R were originally described by Muyzer et al^67^.

#### Growth of ASF361 on GF small intestine mucus

Luminal contents were removed using tweezers from small intestines and mucus was scraped using a frosted glass slide. Mucus was resuspended in sterile water at 1 mg/μL. In a 96-well plate, 10 μL of a 1:1000 dilution of an overnight culture of ASF361 was added to 190 μL of diluted mucus and incubated anaerobically at 37°C using an anaerobic GasPak system. Following incubation, bacterial colonies were enumerated and CFU were calculated based on dilution factor. Bacteria were diluted in sterile 1x PBS and plated in triplicates of 5 μL droplets on MRS agar and incubated anaerobically overnight at 37°C before counting. Bacterial burden was expressed as CFU per well (log(CFU/well).

#### O-and N-glycomic analysis

Approximately 25 mg of tissue was transferred to a 1.5-mL tube and suspended in 1 mL of 50 mM ammonium bicarbonate (ABC). Tissues were probe-sonicated in 15-s intervals separated by 15-s rests, for a total sonication time of 60 s. Samples were centrifuged at 3,000 × *g* for 10 min at 4°C, and the supernatant was transferred to a clean 1.5-mL tube. Protein concentration was measured using a Pierce BCA Protein Assay Kit (part no. 2161296). For each sample, 500 µg of protein was reduced with 5 mM dithiothreitol for 45 min at 50°C. Samples were cooled to 25°C and alkylated with 12 mM iodoacetamide in the dark for 45 min at 25°C. Samples were desalted using 10-kDa molecular-weight-cutoff filters. Each sample was loaded onto a filter and centrifuged at 16,000 × *g* for 10 min. The filter was then washed five times with 350 µL of ABC. To recover the sample, the filter was inverted over a clean 1.5-mL tube and briefly centrifuged, then rinsed with 50 µL of ABC.

Trypsin was added at a 1:20 enzyme-to-protein ratio by mass, and samples were incubated at 37°C for 16 h. Digestion was stopped by heating the samples to 100°C for 1 min. Samples were cooled to 25°C, supplemented with 4 µL of peptide-N-glycosidase F (PNGase F), and incubated at 37°C for 16 h.

Released N-glycans were separated from peptides and O-glycopeptides using C18 solid-phase extraction cartridges. Cartridges were activated with 1 mL of methanol and conditioned with 3 mL of 5% acetic acid in water. Samples were applied to the cartridges, and the flow-through containing released N-glycans was collected. The cartridges were washed with 3 mL of 5% acetic acid. Peptides and O-glycopeptides were then eluted sequentially with 2 mL each of 20% 2-propanol in 5% acetic acid, 40% 2-propanol in 5% acetic acid, and 100% 2-propanol. The N-glycan and peptide/O-glycopeptide fractions were lyophilized and processed separately.

#### N-glycan permethylation and mass spectrometry

Released N-glycans were permethylated using a dimethyl sulfoxide/sodium hydroxide suspension and iodomethane, as described^68^ with modifications. Dried samples were resuspended in 200 µL of dimethyl sulfoxide, followed by 300 µL of dimethyl sulfoxide/sodium hydroxide suspension and 100 µL of iodomethane. Samples were mixed and agitated for 20 min. The addition of dimethyl sulfoxide/sodium hydroxide suspension and iodomethane and the 20-min reaction were then repeated. After the second reaction, samples were quenched with LC–MS-grade water. Permethylated glycans were extracted with dichloromethane, and the organic layer was transferred to a clean tube and dried under a stream of nitrogen.

For matrix-assisted laser desorption/ionization analysis, dried permethylated N-glycans were dissolved in 20 µL of methanol. A 2-µL aliquot was mixed with 2 µL of 15 mg/mL 2,5-dihydroxybenzoic acid prepared in acetonitrile, water, and formic acid at a ratio of 700:300:1 and spotted onto a MALDI target plate. Data were acquired using a Bruker rapifleX MALDI-TOF mass spectrometer. Spectrometer settings used in the RapidfleX were set with an ion source of 20 kV, PIE 2.54 kV, lens 11.8 kV, reflector 1 20.85 kV, reflector 2 1.9 kV, and reflector 3 at 8.6 kV. Mass range detection was set to 1100 to 7000 m/z and peak detection was set to centroid with a signal to noise threshold of 2.0. Raw data was manually interpreted using GlycoWorkbench 2.0.

The remaining samples were dried again under nitrogen and resuspended in 100 µL of methanol. A 30-µL aliquot was mixed with 20 µL of 1 mM lithium acetate, and 20 µL was introduced into an Orbitrap Fusion mass spectrometer using an Ultimate 3000 RSLCnano liquid-chromatography pump connected to an Orbitrap Fusion Lumos MS (ThermoFisher Scientific, Wltham, MA) A commercial nano-LC column (Thermo Scientific DNV PepMAP Neo, 15x0.075 cm; 3 µm packing) was used for chromatography. Separation was carried out in a linear gradient at 50 °C from low to high acetonitrile using Buffer A (aqueous 1mM LiAc in 0.1% [v:v] formic acid) and Buffer B (80% [v:v] acetonitrile with 0.1% [v:v] formic acid in water). The LC flow rate was set to 0.25 µL/min with an initial Buffer B at 5% and linearly increased to 30% at 3 min. Buffer B increased to 60% at 30 min, then to 85% at 50 min. Buffer B was increased to 99% at 60 min and maintained at 99% until 65 min. Buffer B was gradually decreased to 5% at 72 min. The Ion source in the MS was set on positive mode. Precursor ion scans were acquired in the orbitrap at a 120K resolution at a time frame of 3 seconds with a scan range of 500 – 2000. RF lens of 120% was used with absolute ACG of 5.0 x 10^5^, data was collected in profile mode. Precursor ions were selected for subsequent CID fragmentation at 45% collision energy were isolated in the quadrupole with an isolation window of 2 m/z. Fragment ions were analyzed in the Orbitrap with 30K resolution and a scan range set to automatic. Absolute AGC was set to 4.5 x 10^4^ was used with 1 microscan, data was collected in centroid mode.

An initial glycan mass list was generated from the MALDI-TOF data and used to calculate expected [M+Li] ions with charge states from +1 to +5. Candidate masses were then searched in the raw LC–MS data, with the MALDI-TOF results used as a reference for manual interpretation.

#### O-glycan release, permethylation, and mass spectrometry

O-glycans were released from the peptide/O-glycopeptide fraction by reductive β-elimination. Dried samples were incubated with 20 mg of sodium borohydride in 500 µL of 5 mM sodium hydroxide at 50°C for 16 h. Reactions were stopped by slowly adding 10% acetic acid in water. Samples were desalted sequentially using a DOWEX H⁺ resin column and a C18 cartridge and then lyophilized. Residual borate was removed by repeated evaporation with a 9:1 methanol-to-acetic-acid solution under a stream of nitrogen.

Released O-glycans were permethylated using a dimethyl sulfoxide/sodium hydroxide suspension and iodomethane, as described^68^. Dried samples were resuspended in 200 µL of dimethyl sulfoxide, followed by 300 µL of dimethyl sulfoxide/sodium hydroxide suspension and 100 µL of iodomethane. Samples were mixed and agitated for 20 min. The addition of dimethyl sulfoxide/sodium hydroxide suspension and iodomethane and the 20-min reaction were then repeated. Samples were quenched with LC–MS-grade water, and permethylated O-glycans were extracted with dichloromethane. The organic layer was transferred to a clean tube and dried under nitrogen.

Permethylated O-glycans were resuspended in 100 µL of methanol. A 50-µL aliquot was mixed with 50 µL of 1 mM sodium hydroxide, and 20 µL was introduced into an Orbitrap Fusion mass spectrometer using an Ultimate 3000 RSLCnano liquid-chromatography pump. Parameters used for the liquid chromatography and mass spectrometry were the same used in the N-glycan analysis with the exception of Buffer A was composed of aqueous 1mM NaAc in 1% [v:v] formic acid. Mass-spectrometry data were interpreted manually.

#### Glycan quantification and statistical analysis

Peak areas were measured for each identified glycan. The relative abundance of each glycan was calculated as its peak area divided by the sum of all glycan peak areas detected in the corresponding sample. Relative abundances were averaged across biological replicates, and groups were compared using two-tailed unpaired Student’s *t*-tests.

---

**Figure S1.**
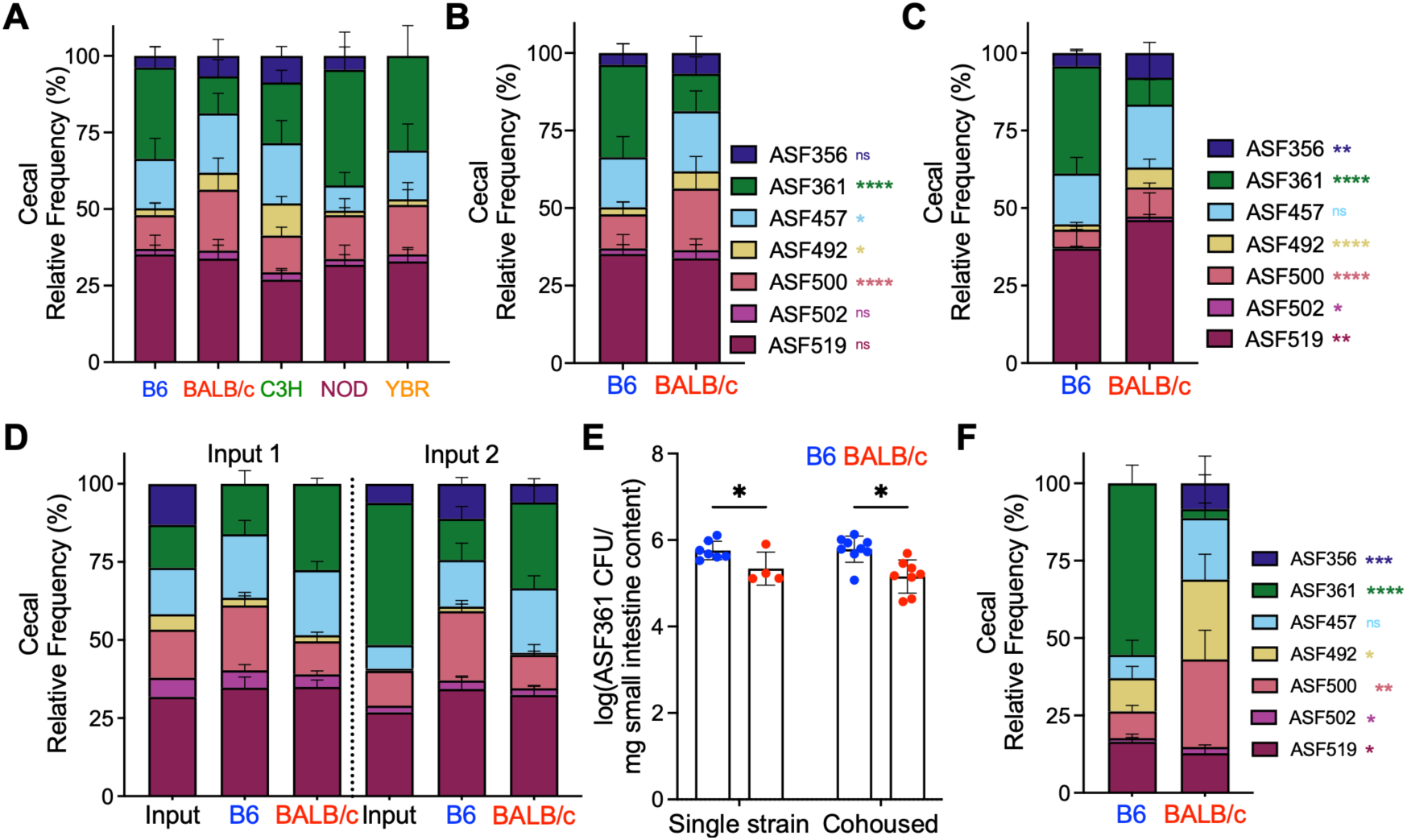
Host genetics shapes *Lactobacillaceae* abundance, related to. Figure 1. (A-B) Total cecal microbiome composition of cecal ASF361 in ASF-colonized mice. (C) Total cecal microbiome composition in 8-week-old adult progeny of ASF-colonized mice. (D) Total composition of two ASF inocula and the cecal microbiome composition of ASF-colonized mice. (E) Small intestine ASF361 CFU in monocolonized mice housed separately by strain or cohoused. (F) Total cecal microbiome composition in mice after 1 week of colonization with ASF. Relative frequency determined by 16S rRNA-seq. Each dot represents an individual mouse. Data were pooled from 2-9 independent experiments. BALB/c, BALB/cJ; C3H, C3H/HeN; B6, C57BL/6J; NOD, NOD/ShiLtJ; YBR, YBR/Ei. Statistical significance was assessed using two-way ANOVA (B-C, F) or a two-tailed unpaired Student’s *t*-test (E). Data are presented as mean ± SD. \**P* < 0.05, \*\**P* < 0.01, \*\*\**P* < 0.001, \*\*\*\**P* < 0.0001; ns, not significant.

**Figure S2.**
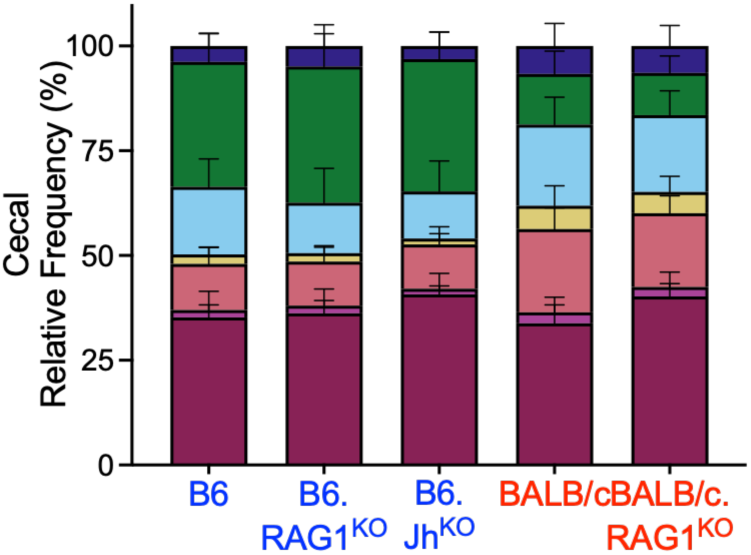
Suppression of ASF361 in BALB/c mice is independent of adaptive immunity, related to. Figure 2. Relative frequency of cecal ASF361 in ASF-colonized RAG1-and JH-deficient mice. Mice were colonized with ASF at 8 weeks of age and analyzed 2 weeks later. Relative frequency determined by 16S rRNA-seq. Data were pooled from 1-9 independent experiments.

**Figure S3.**
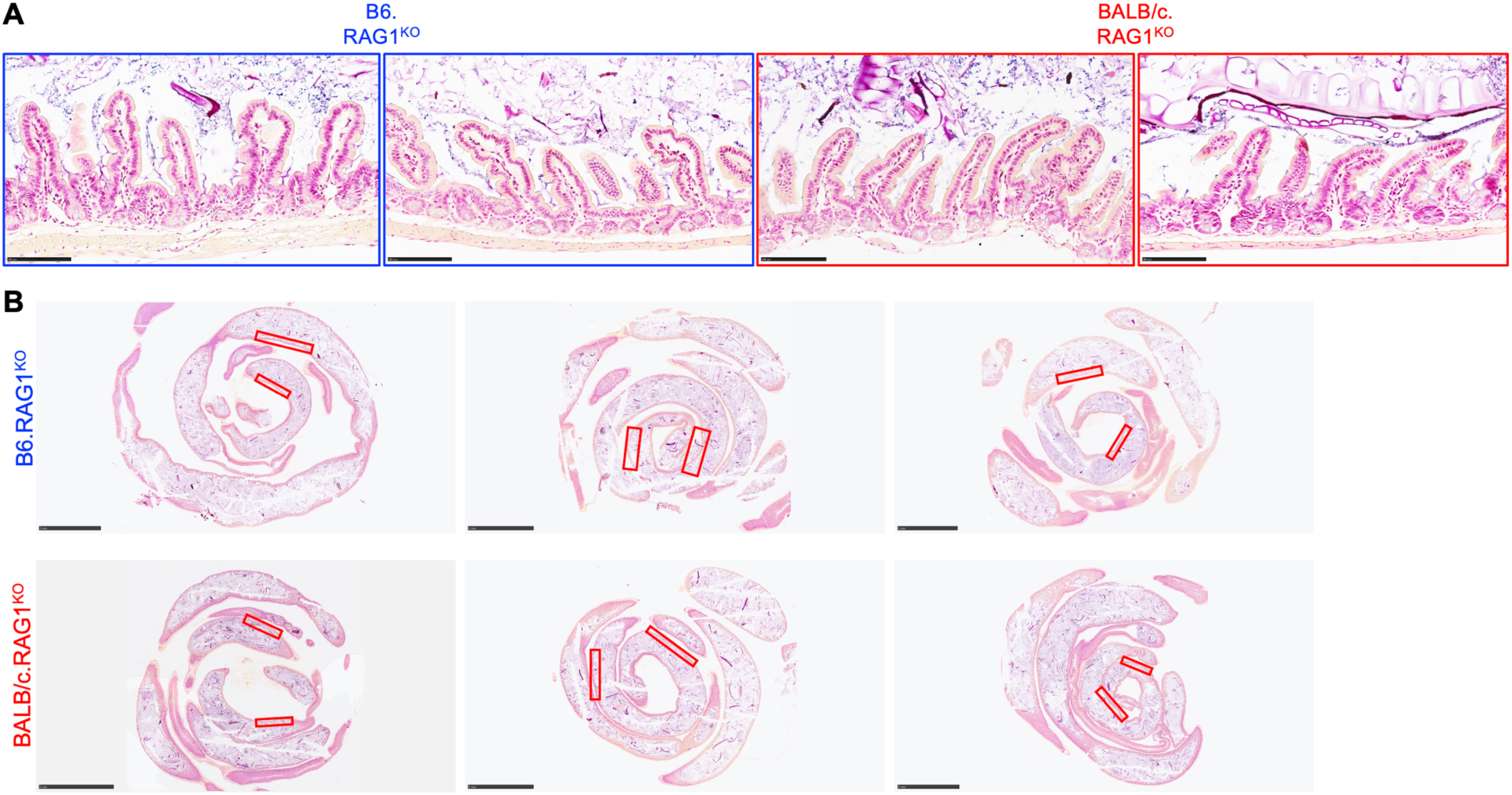
ASF361 is excluded from BALB/c ileal crypts, related to. Figure 3. (A) Gram staining of Carnoy’s-fixed ileal sections from mice monocolonized with ASF361. Each bar is 100 μm. (B) Whole slide scans of Gram-stained Carnoy’s-fixed small intestine tissue from monocolonized B6.RAG1^KO^ and BALB/c.RAG1^KO^. Red rectangles indicate areas that were used for enumeration of bacteria-occupied crypts from Figure 3B. Each bar is 5 mm. Mice were colonized ASF361 at 8 weeks of age and analyzed 2 weeks later.

**Figure S4.**
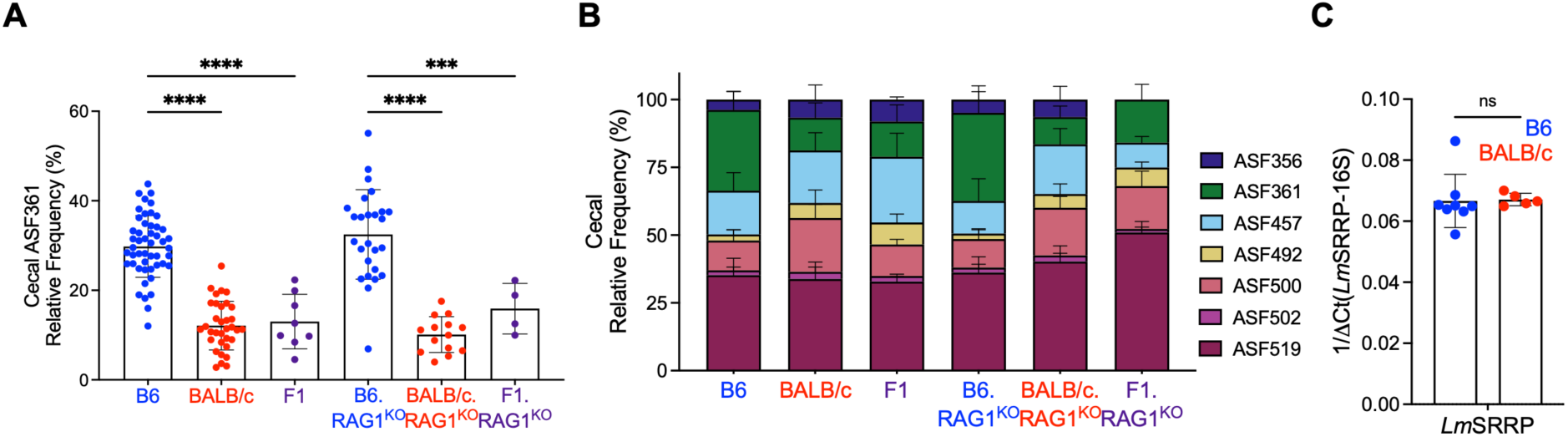
Other bacteria contribute to suppression of ASF361 in BALB/c mice, related to. Figure 4. (A) Relative frequency of cecal ASF361 and (B) total cecal microbiome composition in WT and RAG1-deficient F1 mice colonized with ASF. F1, (B6 x BALB/c)F1; F1.RAG1^KO^, (B6.RAG1^KO^ x BALB/c.RAG1^KO^)F1. F1, (B6 x BALB/c)F1. Mice were colonized with ASF at 8 weeks of age and analyzed 2 weeks later. Relative frequency determined by 16S rRNA-seq. Data were pooled from 2-9 independent experiments. (C) qPCR analysis of *Lm*SRRP expression in ASF361 after 24 hours of growth on small intestine mucus from germ-free mice. Statistical significance was assessed using two-tailed unpaired Student’s *t*-tests (A, C). Data are presented as mean ± SD. \*\*\**P* < 0.001, \*\*\*\**P* < 0.0001; ns, not significant.

